# Identification of diverse antibiotic resistant bacteria in agricultural soil with H_2_^18^O stable isotope probing and metagenomics

**DOI:** 10.1101/2023.01.24.525391

**Authors:** Marcela Hernández, Shamik Roy, C. William Keevil, Marc G. Dumont

**Author notes:** Corresponding author: Marc G. Dumont, University of Southampton, Life Sciences Building 85, Highfield Campus, Southampton SO17 1BJ, UK.

## Abstract

**Background:** In this study, we aimed to identify bacteria able to grow in the presence of several antibiotics including the ultra-broad-spectrum antibiotic meropenem in a British agricultural soil, by combining DNA stable isotope probing (SIP) with high throughput sequencing. Soil was incubated with cefotaxime, meropenem, ciprofloxacin and trimethoprim in ^18^O-water. Metagenomes and the V4 region of the 16S rRNA gene from the labelled “heavy” and the unlabelled “light” SIP fractions were sequenced.

**Results:** After incubations, an increase of the 16S rRNA copy numbers in the “heavy” fractions of the treatments with ^18^O-water compared with their controls was detected. The treatments resulted in differences in the community composition of bacteria. Members of the phyla Acidobacteriota (formally Acidobacteria) are highly abundant after two days of incubation with antibiotics. Several Pseudomonadota (formally Proteobacteria) including *Stenotrophomonas* were prominent after four days of incubation. Furthermore, a metagenome-assembled genome (MAG-1) from the genus *Stenotrophomonas* (90.7% complete) was retrieved from the heavy fraction. Finally, 11 antimicrobial resistance genes (ARGs) were identified in the unbinned-assembled heavy fractions, and 10 ARGs were identified in MAG-1. On the other hand, only two ARGs from the unbinned-assembled light fractions were identified.

**Conclusions:** The results indicate that both non-pathogenic soil-dwelling bacteria as well as potential clinical pathogens are present in this agricultural soil, and several ARGs were identified from the labelled communities, but it is still unclear if horizontal gene transfer between these groups can occur.

## Introduction

Soil represents a natural reservoir of antimicrobial resistance genes (ARG) that has originated as a defence mechanism for the microbes to deter antimicrobial products secreted by competing microbes in the same niche. However, the abuse and/or misuse of antibiotics among humans to treat diseases, and livestock production systems to increase yield has initiated the alteration of natural antimicrobial resistance (AMR) and subsequent spread across all terrestrial ecosystems [1]. Indeed, no soil environment can be now considered pristine as the ARGs are present in garden soils [2], agricultural soils [3], forests [4], grasslands [5], and even Antarctic soils [6]. So much so that soil can harbour up to 32% of the overall ARG diversity [7]. In addition, a previous study has reported the importance of low abundance antibiotic-resistant microbes in soil-plant systems for the spread of AMR [8].

Transmission of AMR back to humans through soil-microbe-animal-plant nexus endangers public health, since the spread of AMR could push us to the pre-antibiotic era. We now know what can happen to AMR in soils due to rise in antibiotic diversity and abundance in the environment but how AMR will spread in soil remains to be seen. In other words, the drivers, or mechanisms of the inevitable spread of AMR in soils when challenged with antibiotics remains to be determined. Deciphering this knowledge gap is crucial for us to develop strategies to alleviate the spread of AMR in terrestrial ecosystems.

It has been hypothesised that the spread of AMR in soil is primarily driven by two non-independent processes that can operate in tandem to alter the antibiotic resistome in soil [1,9]. One process is horizontal gene transfer (HGT) of antimicrobial resistance genes (ARG) between microbial community members. Secondly is the directional selection of antibiotic resistant microbes that can grow in the presence of antibiotics. This could be either due to incorporation of microbiomes derived from anthropogenic sources (e.g., organic fertiliser), or selection and proliferation of naturally resistant microbiota. We are now beginning to understand how HGT can facilitate the spread of AMR in pristine environments [6,10]. For instance, the *bla*_NDM-1_ gene that confers resistance to carbapenem (last resort antibiotic) is now ubiquitous due to successive and distinct HGT events [11,12]. On the other hand, there is limited knowledge about the community composition of the microbiome that can resist antibiotic in soil. One of the main reasons could be the large abundance of extracellular DNA (eDNA) in soil, which cannot distinguish active antibiotic resistant microbes from dead/dormant antibiotic sensitive microbes [13,14]. This could be the reason why studies have reported contradictory results of no change in microbiomes to complete change upon antibiotic addition [5,15].

Agricultural ecosystems represent 38% of the Earth’s ice-free terrestrial surface — the largest use of land on the planet [16]. Sustainable agricultural practice envisions the widespread adoption of organic fertilisers instead of chemical fertilisers as a source of nutrients to maintain or increase crop yield [17]. This is essential to achieve climate-change goals and concomitantly meet the dietary demands of 9 billion people by 2050. However, the build-up of antibiotic concentrations and ARG abundance in environmentally sustainable organic fertilisers, such as livestock manure and sewage sludge, permeates agricultural soils to spread AMR by altering the microbiome [1,9,13,18]. Since AMR microbes are one of the major determinants of AMR spread, it is therefore crucial for us to identify the active fraction of the soil microbial community that can grow in the presence of antibiotics.

Stable isotope probing (SIP) with [^18^O]-water presents a unique approach to identify the active AMR microbes [19,20]. SIP is a cultivation-independent approach that requires the addition of stable-isotope-enriched substrates (e.g., ^13^C-methane, ^18^O-water) to environmental samples followed by analyses of labelled DNA or RNA [20]. SIP techniques can target phylogenetically constrained metabolic processes (e.g., ethane oxidation) where from a diverse pool of active microbial community only those microbial guilds that can assimilate and subsequently incorporate the labelled substrate into their biomolecules such as DNA and RNA are identified. In contrast, SIP-H_2_^18^O as a substrate can potentially label all metabolically active or growing microbes since water is a prerequisite for growth and cellular maintenance [19,21]. Here, fast growing microbes are labelled first, but eventually all active microbes are expected to contain isotope-enriched DNA. Additionally, ^18^O has two more neutrons than naturally abundant ^16^O, whereas ^2^H, ^13^C and ^15^N has only one additional neutron compared to their naturally abundant counterparts (^1^H, ^12^C and ^14^N). This can potentially increase the degree of physical separation of labelled ^18^O-DNA from unlabelled DNA during isopycnic centrifugation in SIP. As a result, the SIP-H_2_^18^O has been used as a robust method to identify the active microbes in a multitude of treatment set-ups such as nutrient addition to soil [22], soil rewetting [21,23], and soil warming [24].

In this study we combine antibiotic selection and SIP-H_2_^18^O to elucidate active and growing microbial communities in an agricultural soil. Antibiotic selection or continuous presence of antibiotics in the experiment will select only AMR microbes to grow, and simultaneously kill or inhibit the growth of sensitive microbes. We also use agricultural soil with no history of antibiotic addition either directly, or indirectly via organic fertilisers. This was done to reduce the bias in identification that can be introduced from long-term exposure of microbes to anthropogenic derived antibiotics as it may already have selected for a resistant microbial community with no difference between the antibiotic challenged and unchallenged communities. Our objectives for this study were to investigate whether microbes in agricultural soil with no-antibiotic history can grow if challenged with antibiotic; secondly, if they are present to identify the microbes and the metabolic machinery and/or AMR genes that confer resistance. We hypothesise that the addition of antibiotics to agricultural soils in the presence of H_2_^18^O will unravel the identity of antibiotic resistant microbes and this will help to understand the drivers of AMR spread.

## Results

To evaluate whether microbes in an agricultural soil with no-antibiotic history can grow if challenged with antibiotic, agricultural soils were incubated with an antibiotic cocktail of meropenem (mem), cefotaxime (ctx), ciprofloxacin (cip) and trimethoprim (tmp), along with H_2_^18^O or natural isotope abundance water (referred to as H_2_^16^O). Here we report the results after 4-days of incubation with antibiotic addition at 0 and 48 hr time-points. A total of 18 CsCl gradient fractions were collected following ultracentrifugation and the 16S rRNA gene copy numbers (bacterial abundance) were analysed for each experimental setup.

Incubation with H_2_^18^O increased the overall buoyant density of the extracted DNA as compared to the H_2_^16^O controls (Figure 1). ^18^O-labelled DNA (heavy fractions) resided in fractions with densities 1.73 g ml^-1^ and above, whereas unlabelled DNA (light fractions) resided in fractions with densities 1.729 g ml^-1^ and lower (Figure 1). This indicates that bacteria were actively incorporating ^18^O into their DNA. Here, the heavy fraction indicates active or growing microbes, whereas the light fraction indicates dormant or dead microbes.

**Figure 1.**
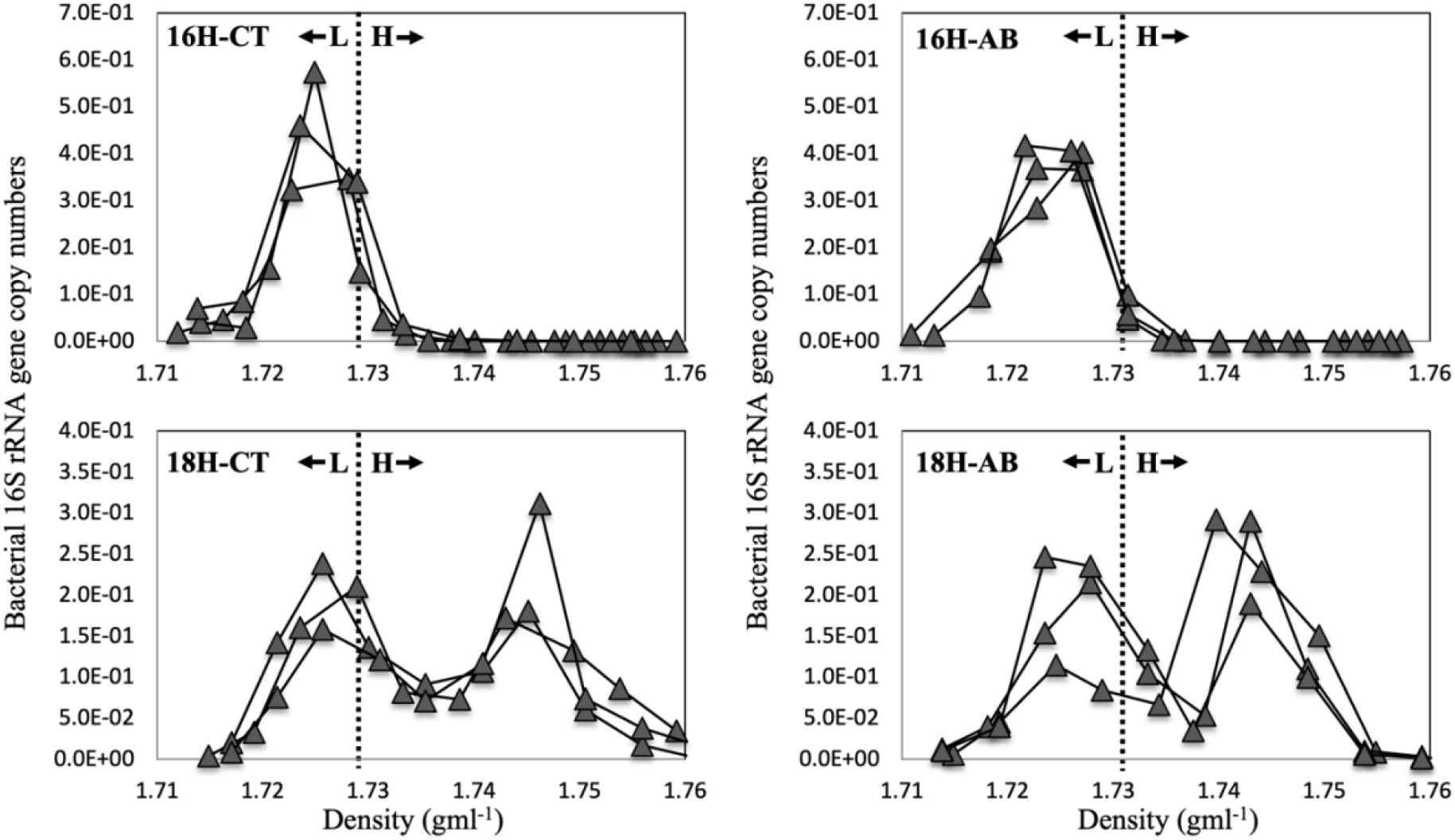
Abundance of bacterial 16S rRNA genes in CsCl density gradients after ^16^O-/^18^O-H_2_O incubation. Vertical dotted lines demarcate the heavy (H) fractions from light (L) fractions. Each line represents a sample. 16H-CT: soils incubated in the presence of natural isotope abundance water (H_2_^16^O) without antibiotics (control); 16H-AB: incubation in the presence of H2^16^O and antibiotics; 18H-CT: incubation in the presence of H_2_^18^O without antibiotics (control); 18H-AB: incubation in the presence of H_2_^18^O and antibiotics.

After 4-days of incubation with H_2_^18^O there was a large abundance of bacterial 16S rRNA gene copies in the heavy fraction as compared to the heavy fraction of samples incubated with H_2_^16^O (Figure 1). This was the case for both the antibiotic treatment and no-antibiotic controls suggesting there was substantial bacterial growth in the presence of antibiotics.

Species richness was significantly lower in the heavy fractions (2278 ± 142, mean ± 95% confidence interval) than light fractions (5025 ± 99) for antibiotics (AB) treated soil (p<0.001). Similarly for CT treatment (i.e., control: soil without antibiotics), the species richness was significantly lower (p=0.013) in the heavy fractions (2926 ± 314) than in the light fractions (3980 ± 370). Shannon diversity (*H*) was lower (p<0.001) for heavy fractions (1.99 ± 0.36) than light fractions (6.50 ± 0.37) for AB treated soil. However, for CT, Shannon diversity (*H*) did not differ (p=0.157) between heavy fractions (4.37 ± 0.12) and light fractions (5.12 ± 0.68). Evenness (*J*) indexes were lower (p<0.001) in heavy fractions (0.42 ± 0.07) than in light fractions (0.98 ± 0.01) for AB treated soil. Contrarily, for control treatments (CT), evenness (*J*) did not differ (p=0.219) between heavy fractions (0.93 ± 0.01) and light fractions (0.93 ± 0.05) (Figure 2).

**Figure 2.**
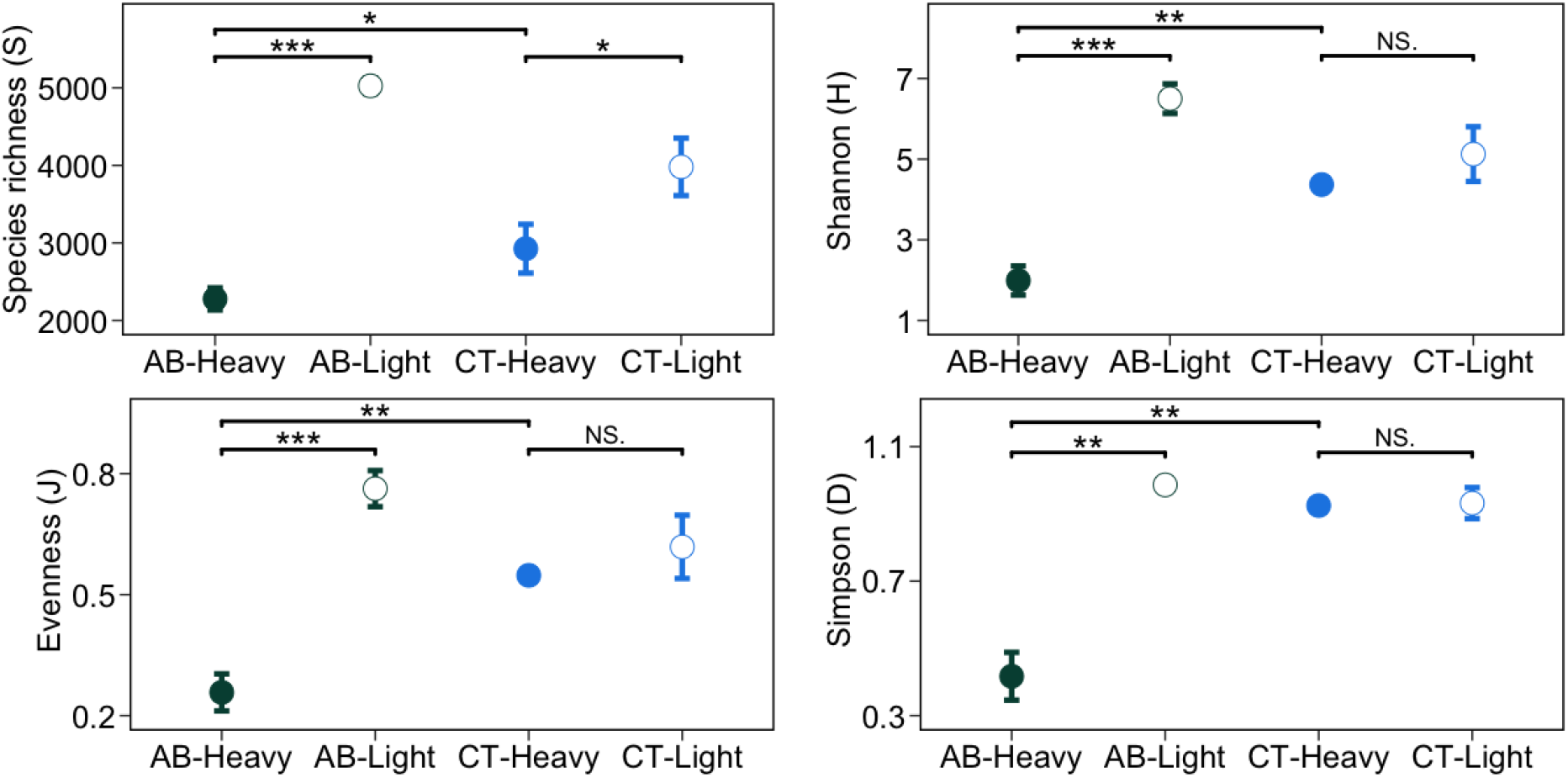
Alpha-diversity of 16S rRNA gene sequences from the “heavy” and “light” fractions of DNA extracted from soils incubated with H_2_^18^O in the presence (AB) or absence (CT) of antibiotics. Alpha-diversity is summarised as species richness (S), Shannon diversity (H), evenness (J), and Simpson (D). “*” corresponds to p-value <0.05 from pairwise t-test; “**” corresponds to p-value <0.01; “***” corresponds to p-value <0.001; “NS.” corresponds to p-value >0.05

When comparing the heavy fractions of AB and CT treatments, species richness was lower (p=0.039) for heavy fractions for AB treatments (2278 ± 142) than CT treatments (2926 ± 314). Similarly, Shannon diversity was lower (p=0.003) for AB treatments (1.99 ± 0.36) than CT treatments (4.37 ± 0.12). Evenness was also lower (p=0.005) for AB treatments (0.42 ± 0.07) than CT treatments (0.93 ± 0.01) (Figure 2). Finally, the coefficient of variation (CV) for all alpha diversity indices across all the treatments ranged from 0.4% to 16.0% (Figure 2).

The microbial community composition was consistent for all replicates of both heavy and light fractions across all the treatments. The community composition for heavy and light fractions of AB and CT when incubated with H_2_^18^O were different as they clustered separately (Figure 3). Community composition of the light fraction in H_2_^18^O incubated CT soil (18H-CT-Light) was similar to both heavy (16H-CT-Heavy) and light (16H-CT-Light) fraction in H_2_^16^O incubated CT soils as shown by their proximity in the PCoA plot and the relative abundance profile (Figure 4, S3). Together, these three fractions (18H-CT-Light, 16H-CT-Heavy, 16H-CT-Light) along with light fraction of H_2_^16^O incubated AB soil (16H-AB-Heavy) were similar to the composition of the original soil.

**Figure 3.**
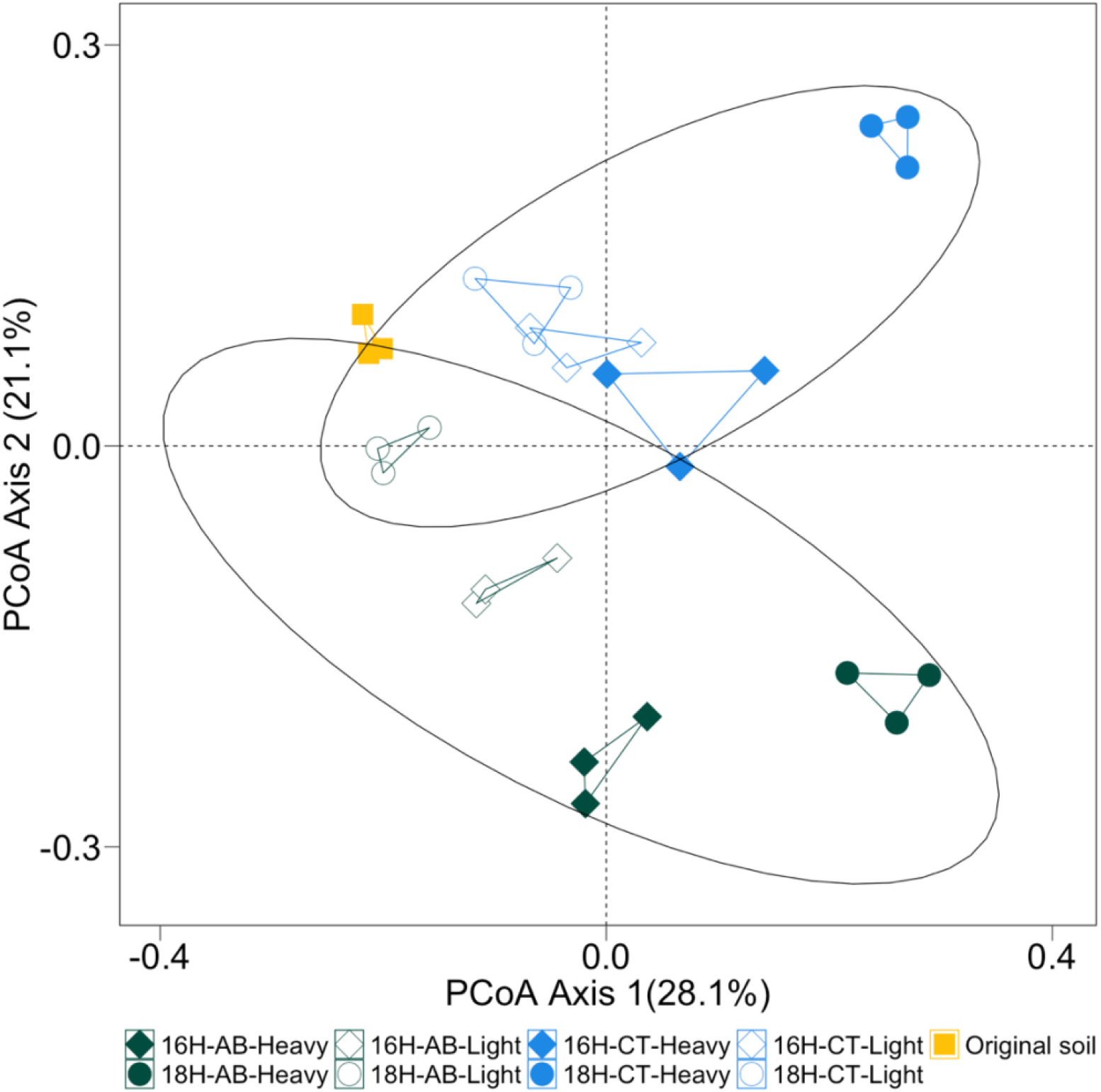
Principal coordinate analysis (PCoA) plots of bacterial OTUs (97% sequence similarity) derived from 16S rDNA extracted from soil. The legend indicates the origin of the samples. 16H: incubation with H_2_^16^O; 18H: incubation with H_2_^18^O; AB: incubation with antibiotics; CT: incubation without antibiotics; Heavy: “heavy” fractions of the extracted soil DNA; Light: “light” fractions of the extracted soil DNA.

**Figure 4.**
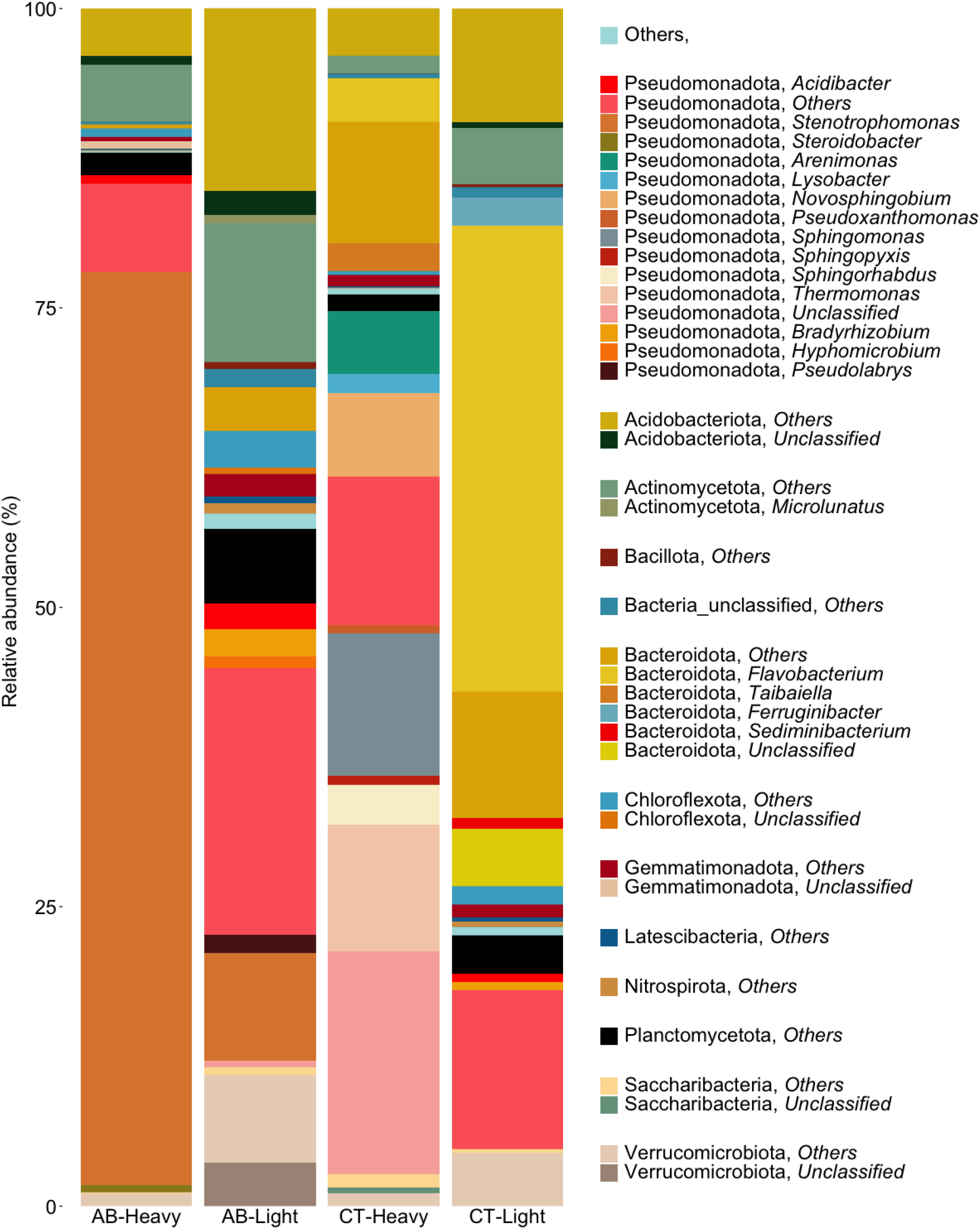
Relative abundance of microbial communities at the phylum level identified in the “heavy” and “light” fractions of DNA extracted from soils incubated with H_2_^18^O in the presence (AB) or absence (CT) of antibiotics.

The community in the heavy fractions of AB incubated with H_2_^18^O were dominated by high relative abundances of Pseudomonadota (84.9 ± 2.86%), Actinomycetota (formally Actinobacteria, 4.8 ± 1.42%), Acidobacteriota (4.7 ± 0.93%), Planctomycetota (formally Planctomycetes, 1.9 ± 0.42%), Verrucomicrobiota (formally Verrucomicrobia, 1.2%), and Gemmatimonadota (formally Gemmatimonadetes, 1.1 ± 0.10%). *Stenotrophomonas* (Pseudomonadota, 76 ± 4.67%) was the most abundant genus (Figure 4). In contrast, the heavy fractions for CT treatments were dominated mainly by Pseudomonadota (72.1 ± 7.9%), Bacteroidota (formally Bacteroidetes, 16.1 ± 5.03%), Acidobacteriota (3.9 ± 0.83%), Saccharibacteria (2.3 ± 2.22 %), Actinomycetota (1.5 ± 0.38%), Planctomycetota (1.3 ± 0.15%), Verrucomicrobiota (1.24%), and Gemmatimonadota (1.1%). Here, *unclassified* (Pseudomonadota, 17.18%), *Sphingomonas* (Pseudomonadota, 11.9%), *Thermomonas* (Pseudomonadota, 10.8%), *Arenimonas* (Pseudomonadota, 5.28%), *Novosphingobium* (Pseudomonadota, 7.03%) were the abundant genera (Figure 4).

The relative abundance in the light fractions of AB were dominated by Pseudomonadota (38.7 ± 5.38%), Acidobacteriota (17.2 ± 0.62%), Actinomycetota (12.3 ± 4.24%), Verrucomicrobiota (10.9 ± 1.01%), Planctomycetota (6.2 ± 1.17%), Bacteroidota (3.7 ± 0.29%), Chloroflexota (formally Chloroflexi, 3.6 ± 0.68%), and Gemmatimonadota (1.9 ± 0.19%). The most abundant genera were *Stenotrophomonas* (Pseudomonadota, 8.9%), *Bradyrhizobium* (Pseudomonadota, 2.3%, only in one replicate), and *Acidibacter* (Pseudomonadota, 2.1%) (Figure 4). In contrast, the light fractions of CT were dominated by Bacteroidota (57.5 ± 8.54%), Pseudomonadota (14.7 ± 2.46%), Acidobacteriota (9.9 ± 2.59%), Actinomycetota (4.7 ± 1.79%), Verrucomicrobiota (4.4 ± 1.29%), Planctomycetota (3.1 ± 0.62%), Chloroflexota (1.52 ± 0.40%), and Gemmatimonadota (1.1 ± 0.12%). The most abundant genus in this treatment was *Flavobacterium* (Pseudomonadota, 38.91 ± 4.67%) (Figure 4).

A heatmap was created to visualise and compare the abundance of the 20 OTUs that explains the most variation in the axis-1 and axis-2 of the PCA ordination. Out of a total of 28 OTUs selected, 19 OTUs belong to Pseudomonadota, followed by six OTUs of Bacteroidota, two OTUs of Verrucomicrobiota, and one OTU of Acidobacteriota. *Stenotrophomonas* (Pseudomonadota; OTU-7) was dominant in the heavy fraction of AB compared to heavy fraction of CT. On the other hand, *Sphingomonas* (Pseudomonadota; OTU-1065, OTU-1321, OTU-2509, OTU-488405, OTU-692415), *Lysobacter* (Pseudomonadota; OTU-12766), *Novosphingobium* (Pseudomonadota; OTU-14845), Xanthomonadaceae (Pseudomonadota; OTU-13089), *Arenimonas* (Pseudomonadota; OTU-1764) were dominant in heavy fraction of CT compared to AB. Additionally, *Pseudolabrys* (Pseudomonadota; OTU-1764), DA101 (Verrucomicrobiota; OTU-424), OPB35 (Verrucomicrobiota; OTU-8196) were dominant in light fractions of AB compared to heavy fractions of AB (Figure 5).

**Figure 5.**
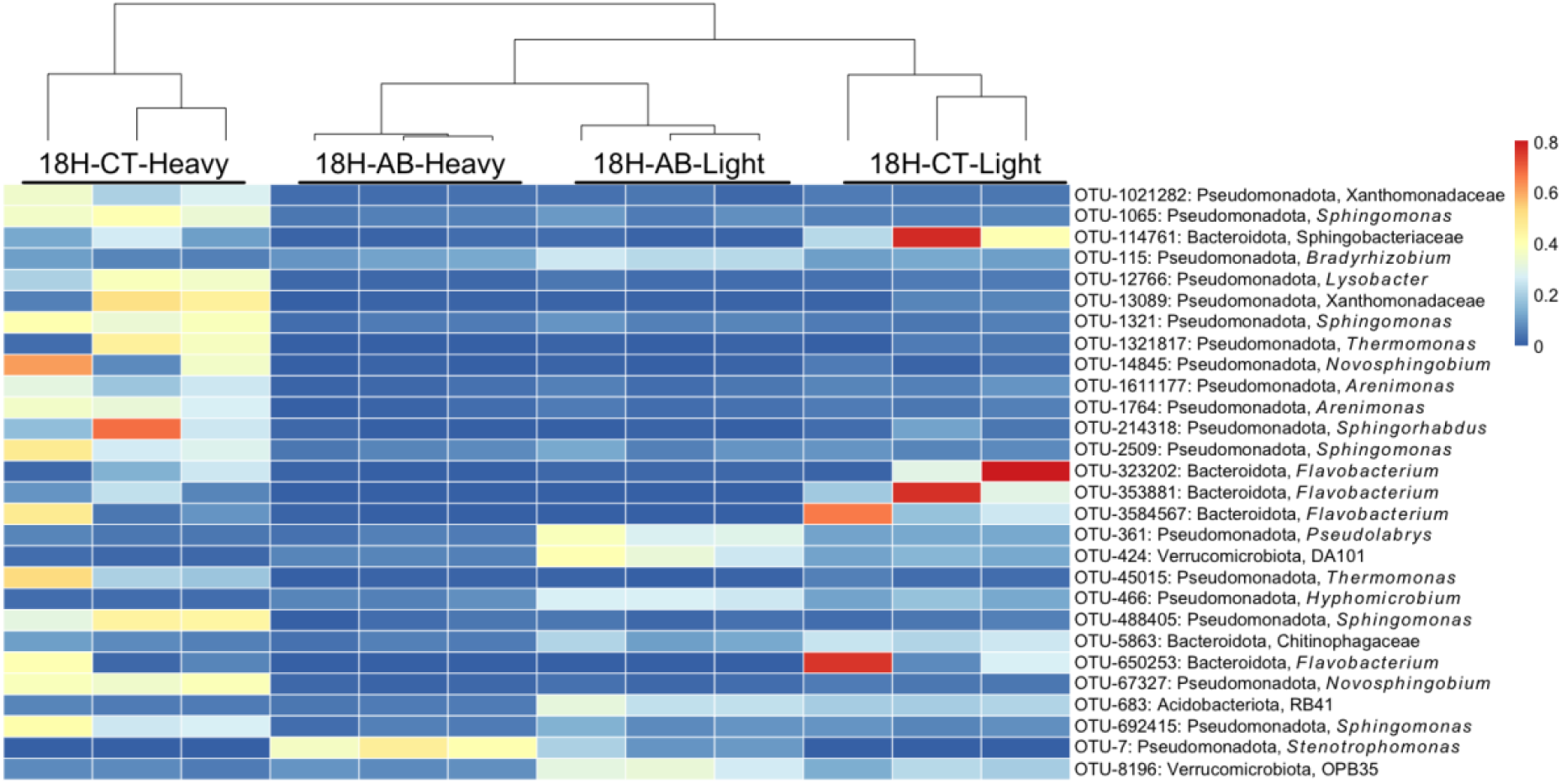
Heatmap of the most relevant bacterial OTUs identified in the “heavy” and “light” fractions of DNA extracted from soils incubated with H_2_^18^O in the presence (AB) or absence (CT) of antibiotics. 18H-AB-heavy indicates “heavy” fractions treated with antibiotic;18H-AB-light indicates “light” fractions treated with antibiotic; 18H-CT-heavy indicates “heavy” fractions without antibiotic treatment; 18H-CT-light indicates “light” fractions without antibiotic treatment. The coloured scale represents the relative abundance of OTUs.

The DNA of both heavy and light fractions of soil when incubated with ^18^O-labelled water in the presence of antibiotics were sequenced individually using high-throughput sequencing. After genome binning of both heavy and light fractions, one qualified metagenome-assembled genome (MAG) was generated with 90.7% completeness and 0% contamination. The MAG was affiliated to *Stenotrophomonas* (Pseudomonadota) (Figure 6). This is in-sync with the results of 16S rRNA gene sequencing that also showed *Stenotrophomonas* (Pseudomonadota) as the dominant genus when incubated with antibiotics (Figures 4, 5).

**Figure 6.**
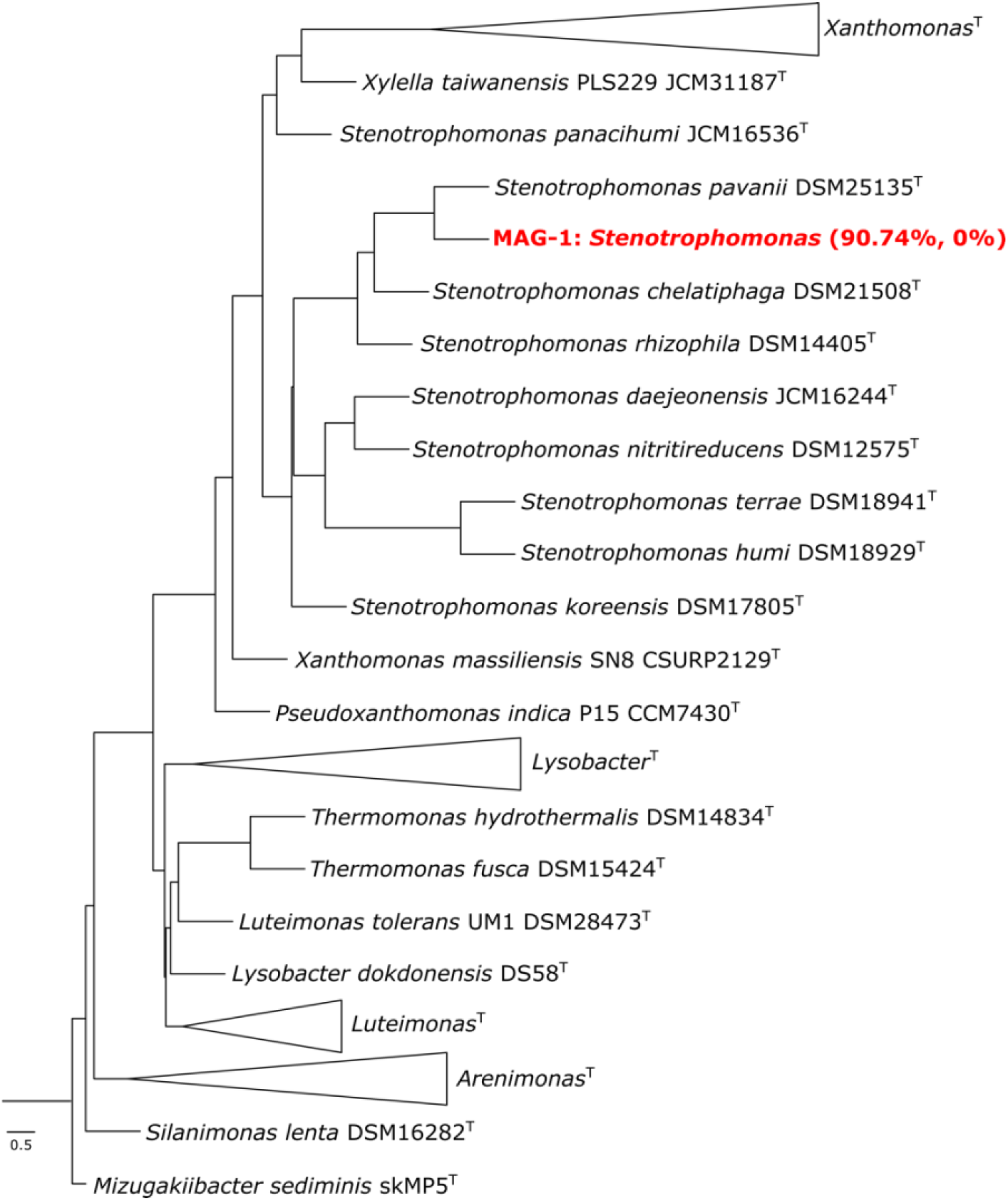
Multi-locus phylogenetic tree of the MAG-1 using autoMLST. 98 conserved housekeeping genes were used for the analyses. MAG-1 is indicated in red together with their respective completeness and contamination.

Analysis of the unbinned-assembled genomes of light and heavy fractions, along with the genome of MAG-1 helped understand whether the microbial community exposed to antibiotic contained antimicrobial resistance genes (ARGs) to survive the stress of antibiotics. The presence of *aph*(*3*’)-*IIc*, with 85.46% similarity to *Stenotrophomonas maltophilia* strain K279a was observed in the heavy fraction and MAG-1, and no presence of this gene was observed in the light fraction. *aph*(*3*’)-*IIc* encodes for the aminoglycoside phosphotransferase enzyme that confers resistance to antibiotics in the aminoglycoside class (butirosin, paromycin, kanamycin, neomycin) among others. Similarly, the presence of *oqxB*, with 76.28% similarity to *Escherichia coli* plasmid pOLA52 in heavy fraction and MAG-1, which was also absent in the light fraction was also observed. The *oqxB* gene encodes for an efflux pump that confers resistance to amphenicol class antibiotics (e.g., chloramphenicol), disinfectants (e.g., benzalkonium chloride, cetylpyridinium chloride), quinolone class antibiotics (e.g., ciprofloxacin, nalidixic acid), trimethoprim, and others. However, *dfrB3*, which encodes for dihydrofolate reductase that confers resistance to trimethoprim, was found with a 90.14% similarity with the plasmid R751 in *Klebsiella aerogenes* only in the light fraction. The presence of ARGs that confer resistance to beta-lactam were also found in both light and heavy fraction, but not in MAG-1. For example, *blaTEM-181* in the light fraction was found with a 99.86% similarity with a vector pUC-3GLA, and *blaL1* in heavy fraction with 85.84% of similarity with a beta-lactamase gene in *Stenotrophomonas maltophilia* strain K1. No ARG conferring beta-lactam resistance was present in MAG-1 (Table 1).

**Table 1.**
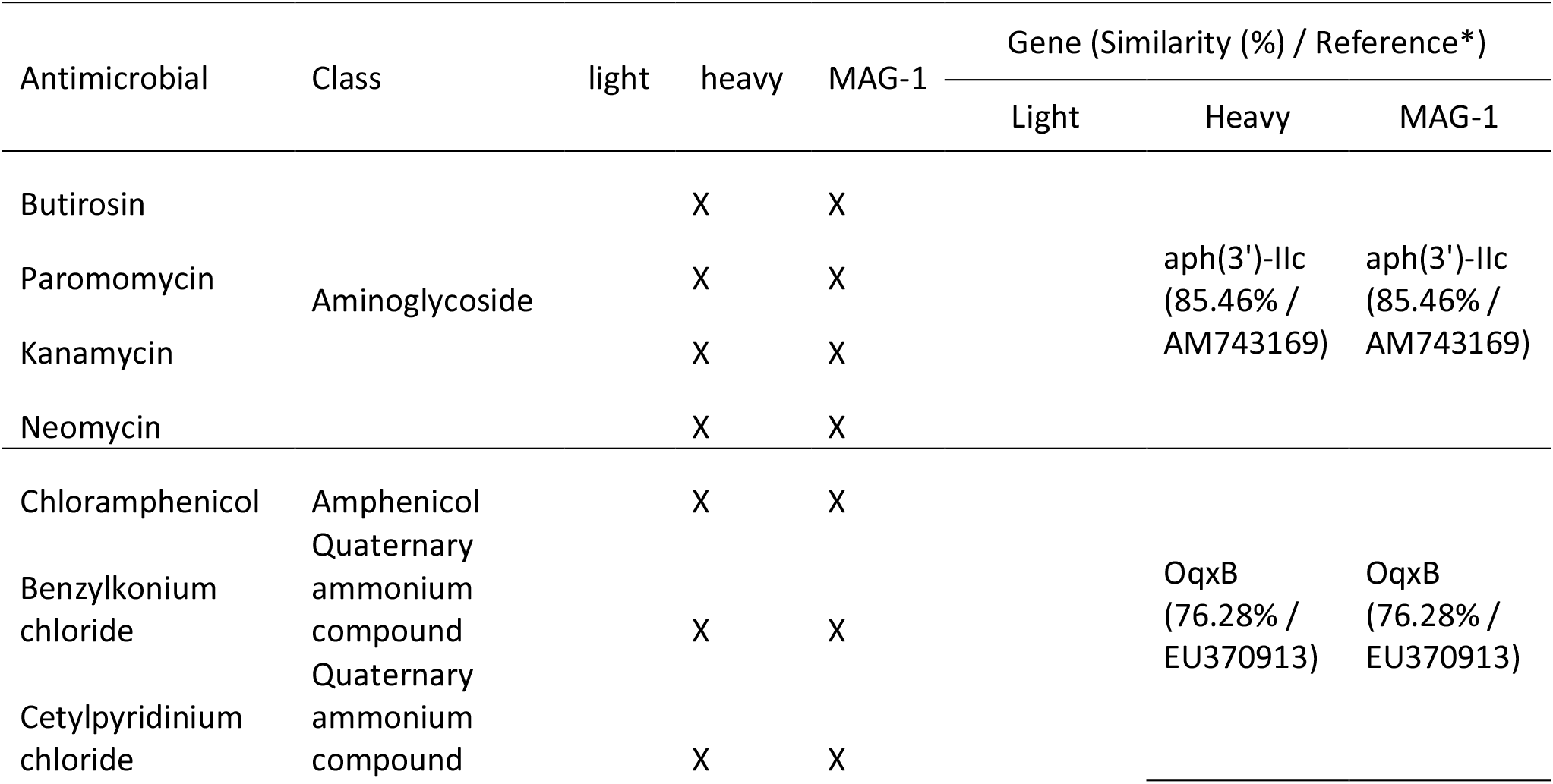
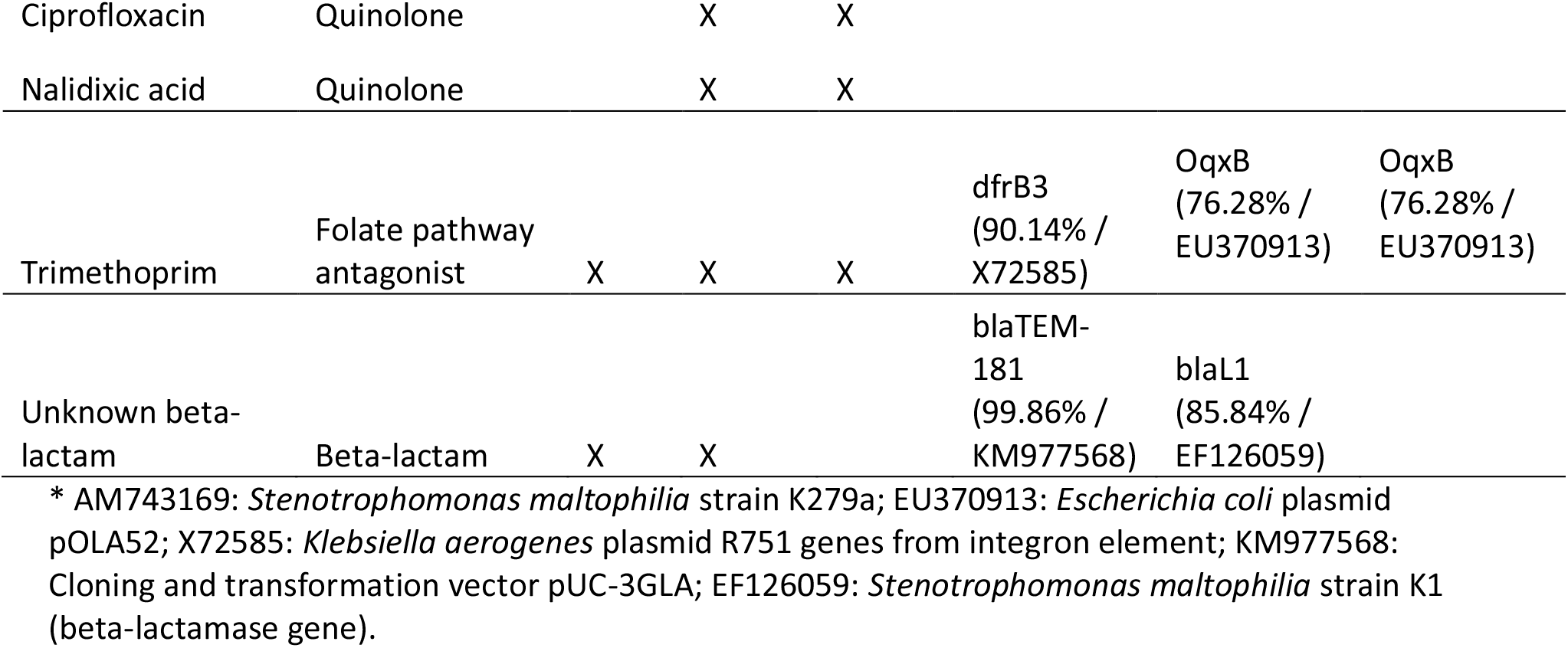
Antimicrobial resistance genes found in the unbinned-assembled reads (from heavy and light fractions) and MAG-1

## Discussion

In this study we used DNA-SIP with H_2_^18^O to identify the antibiotic resistant microbes from the active pool of an agricultural soil microbiome that was not previously exposed to antibiotics. The results showed that microbes can grow in the presence of antibiotic even in the agricultural soil with no-antibiotic history (Figure 1). On the other hand, not all active microbes are antibiotic resistant, since community composition was different between antibiotic treated and untreated soil (Figures 2–5). The metagenomic analyses revealed the presence of ARGs in the active resistant microbial community (Table 1). Additionally, a MAG belonging to *Stenotrophomonas* was found in the heavy fraction after incubation with H_2_^18^O and antibiotics. The study highlights the ability of DNA-SIP with H_2_^18^O to identify active antibiotic resistant microbes.

The results showed that microbes in soils without prior exposure to antibiotic can harbour microbes that can become active and enriched in the presence of antibiotics during a short period of time, in this case after four days of incubation. This suggests that soils contain a seedbank of antibiotic resistance, and the microbiome can shift dramatically towards an enrichment of antibiotic resistant populations even after a short exposure to antibiotics. This also highlights the potential of soil to harbour native AMR bacteria, for these microbes to become dominant, and subsequently spread after exposure to antibiotics. The long-term consequence of shifts in community composition, for example biogeochemical transformations, soil fertility, and disease risk is not clear [25].

The AMR bacterial seedbank in soil could be a result of *in situ* selection as a consequence of natural production of antimicrobials produced by the microbial community as microbes compete for resources. Indeed, soils intrinsically harbour AMR bacteria and are a natural reservoir for ARGs [7,12]. Alternatively, antibiotic resistant bacteria or ARGs could have been introduced to the soil from external sources. This is common in soils exposed to livestock manure or sludge [9,18]. Moreover, the dispersal through unconventional sources such as birds can provide the initial seed for the microbial communities to spread AMR. Birds have been shown to spread AMR through long-distance and localised migration [26,27]. For example, Franklin’s gulls (*Leucophaeus pipixcan*) in Chile were found to have twice the prevalence of ESBL-producing *E. coli* compared to humans in the same area along with high sequence similarity suggesting transmission. Interestingly, the gulls also share sequences with drug-resistant human pathogens identified in clinical isolates from the central Canadian region, which is a nesting place for these gulls [27]. However, more studies are needed to decisively establish the roles of birds to encounter and acquire active antibiotic resistant microbes in soils without prior exposure to antibiotic. Horizontal gene transfer (HGT) is another mechanism for ARG accumulation within a community but there was no evidence for this in the present study.

The active pool of antibiotic resistant microbes was dominated by Pseudomonadota, Actinomycetota, Acidobacteriota (Figures 4, 5). Pseudomonadota are known to be physiologically and metabolically versatile with variable morphology that allows them to subsist in various ecological niches [28–33]. This could be the reason why 72-84% of the OTUs labelled in the presence of antibiotics were affiliated to Pseudomonadota (Figures 4, 5, S3, S4). Further, due to their versatility, Pseudomonadota also contain the greatest number of bacterial pathogens to an extent that this phylum has been proposed to be a potential diagnostic signature for disease risk [34,35]. Actinomycetota is another near ubiquitous phylum in soil that are known for their ability to synthesize diverse secondary metabolites and harbour different ARGs [33,36,37]. It is hypothesised that in soils, ARGs of pathogenic Pseudomonadota originated from Actinomycetota through horizontal gene transfer using conjugative plasmids [38–40]. These results reaffirm the role of Actinomycetota in AMR spread and high abundance in AMR microbiomes. Similarly, Acidobacteriota is also widespread in soil with phylogenetic depth and ecological importance comparable to Pseudomonadota [33,41]. Acidobacteriota can harbour multiple integrative and conjugative elements in their genome, a major determinant of horizontal gene transfer, that confers them a major advantage to survive, resist, and persist in the presence of antibiotic [42,43].

In this study, *Stenotrophomonas* was found to be the dominant genus with a relative abundance of 76% in the active resistant microbiome (Figures 4, 5, 6) and it possessed ARGs for diverse antibiotics (see MAG-1 in Table 1). *Stenotrophomonas* is an antibiotic resistant opportunistic pathogen that is commonly linked to respiratory infections in humans [44]. Possession of a wide range of ARGs by *Stenotrophomonas* in an antibiotic unexposed soil is disturbing, but not unusual and rare. For instance, on one hand, ARG in *Stenotrophomonas* strains has been reported from deep-sea invertebrates [45]. On the other hand, multi-drug resistant *Stenotrophomonas* is a common nosocomial and community-acquired infection [44].

The active pool of antibiotic resistant microbiota in this agricultural soil contains ARGs for a wide variety of antibiotics (Table 1). Surprisingly, many of these antibiotics such as aminoglycosides, chloramphenicol, were not even part of the experiment in this study. This highlights the potential role of the seedbank of resistant bacteria in AMR spread. We hypothesize that these microbes are present in soils at low abundance but with selection can become enriched increasing the probability of causing disease outbreaks in livestock and human populations. Their enrichment may also spread resistance within the microbial community through HGT.

## Conclusion

In this study, the active resistant soil microbiome from an agricultural field with no prior history of antibiotic exposure using DNA-SIP with H_2_^18^O was identified and differences in the composition of active soil microbes and active antibiotic resistant soil microbes was observed. The metagenome data shed light on the diversity of antibiotic resistant genes of the resistant microbiome. In this study, we identified the prevalence of antibiotic resistant *Stenotrophomonas* in the soil, which might be consequential for AMR spread and disease risk. Overall, this study makes a strong case for DNA-SIP with H_2_^18^O to identify the clinically important drug-resistant microbes in the environment. Finally, this method can become gold standard to understand and identify the drivers of AMR spread in any environment.

## Materials and methods

### Soil sampling

Agricultural soils from Chilworth Manor Experimental plots located in the Victorian Walled Gardens (Southampton, U.K.) was sampled in October 2016. This soil does not have history of manure and no antibiotic applications for over 20 years. Also, the last application of herbicide Round-up (Glyphosate at a concentration of 360 g L^-1^ as active ingredient) was in 2007-2008 (M. Cotton, *pers. comm.*). Samples were collected from 10-cm deep in a 10m triangular pattern as described previously (Gomez-Alvarez et al., 2007). In total, three soil samples (not pooled) were transported to the University of Southampton and storage at 4°C for further experiments. Physico-chemical parameters were carried out at the Anglian soil analyses company (Lincolnshire, U.K.) and detailed in Table S1. This soil is a sandy/loam soil with a pH of 6.17 (±0.006), organic matter of 7.73% (±1.43) and dry matter of 85.99% (±3.58).

### Soil incubations

Initial tests were performed to determine the concentration of antibiotic necessary to inhibit bacterial growth in soil for up to 12 days. This is necessary due to potential attenuation of the antibiotic by soil (for methodology see Appendix S1). Since the attenuation of the antibiotic is as fast as two days (Figure S1), a second preliminary experiment was carried out by incubating the soils with several antibiotics to determine the suitable ones to be used for further labelling experiment. Antibiotics were chosen because of their mechanism of action and described in Appendix S1. The resistance genes for these antibiotics have been found in the genome of *Klebsiella pneumoniae* [46]. After performing the preliminary experiments, we decided to incubate the soils for up to 4 days due to its fast decomposition (Figure S1) and four antibiotics were chosen for further incubations with H_2_^18^O (Table S2). One g of soil was incubated in 1.5 ml of with either label water (H_2_^18^O) or unlabelled water (H2O). Antibiotics [meropenem (mem), cefotaxime (ctx), ciprofloxacin (cip) and trimethoprim (tmp)] at a concentration of 100 μg/ml each were added to the slurry at time 0 h and 48 h. Incubations were performed at 200 rpm, dark and room temperature. Sampling time for the treatment was at day 2 and day 4. For controls, sampling was carried out after 4 days of incubation. The experiments were performed in triplicate to provide statistically testable data (Figure S2).

### DNA extraction

DNA was extracted from the soil at the end of the treatments by using the Power-Soil DNA isolation kit (Mo Bio, UK) according to the manufacturer’s recommendation. DNA purity and quantification were determined using a NanoDrop^®^ Spectrophotometer ND-1000 (Thermo Fisher Scientific, USA). All DNA samples were stored at −80°C for further analysis.

### H_2_^18^O-SIP procedure

A standard DNA-SIP protocol was used to resolve [^18^O]-incorporation based on buoyant density [47]. 1 μg of genomic DNA was loaded into 5.6-ml polyallomer quick-seal centrifuge tubes (Beckman Coulter, USA) containing gradient buffer [20] and CsCl. The isopycnic centrifugation of DNA was performed with an initial CsCl buoyant density of 1.725 g mL^-1^ subjected to centrifugation at 177,000 × g for 36-40 h at 20 °C in an Optima XPN-80 ultracentrifuge (Beckman Coulter, USA). At the end of the centrifugation, 18 fractions were separated from each gradient.

### Quantitative PCR

The 16S rRNA gene was quantified in each of the fractions. All qPCR reactions were performed on a StepOne Plus real-time PCR system (Applied Biosystems) and the data were processed using StepOne software v2.3 (Applied Biosystems). For all assays, standards were prepared by PCR of cloned genes. Standards were serially (10^1^–10^7^) diluted and used for the calibration curves in each assay. Controls were run with water instead of template DNA. The assays were based on dual-labelled probes using the primer–probe sets: BAC338F/BAC516P/BAC805R [48]. The probe was synthesized with 6-Carboxyfluorescein (6-FAM) on their 5’end and Black Hole Quencher 1 (BHQ1) on their 3’end. Each reaction was 20 mL in volume and contained the following mixture: 10 μL of TaqMan fast advanced master mix (1X) (Applied Biosystems), 1.0 μL of of primer mix [18 μL BAC338F (0.9 μM), 18 μL BAC805R (0.9 μM), 5 μL BAC516P (0.25 μM) and 59 μL of TE buffer], DNA template (2.0 μL) and 7.0 μL of water. The program used was 95°C for 5 min, followed by 35 cycles of 95°C for 30 s and 62°C for 60 s for annealing, extension and signal acquisition respectively [49]. Efficiencies of 97 to 103% with R^2^ values > 0.98 were obtained.

### High-throughput sequencing

The 16S rRNA genes from SIP gradient fractions was amplified and sequenced by barcoded Illumina sequencing. PCR primers 515FB (GTGYCAGCMGCCGCGGTAA) and 806RB (GGACTACNVGGGTWTCTAAT) from the Earth Microbiome project (http://press.igsb.anl.gov/earthmicrobiome/) targeting the V4 region of the 16S rRNA gene (approximately 250 nucleotides) were used. Library preparation and sequencing was performed at the National Oceanographic Centre (NOC) of the University of Southampton, UK, following methodologies described by [50]. Samples were pooled in an equimolar concentration and sequenced on separate runs for MiSeq using a 2 bp × 300 bp paired end protocol.

The total metagenomic DNA of the heavy and light fractions from incubations with H_2_^18^O (total of six samples) were sequenced on an Illumina MiSeq at the University of Southampton (as described above). The metagenome was analysed on a high-performance computing cluster supported by the Research and Specialist Computing Support Service at the University of East Anglia (Norwich, UK).

### Bioinformatic Analyses

For the 16S rRNA-sequencing, quality filtering of the sequences was carried out by using cutadapt [51]. Forward and reverse reads were then merged by using the usearch fastq_mergepairs command [52]. Downstream processing was performed by using UPARSE [52] and UCHIME pipelines [53]. Briefly, sequences shorter than 250 bp were discarded, singletons were retained, and operational taxonomic units (OTUs) were defined at a sequence identity level of 97%.

For the DNA sequences, reads were checked using FastQC version 0.11.8 [54]. Low-quality reads were discarded using BBDuk version 38.68 [55]. Afterwards, reads were merged into scaffolds using de novo assembler metaSPAdes version 3.13.1 [56]. Binning of the assembled scaffolds from both heavy and light fractions was carried out with the metaWRAP version 1.2.1 [57]. Completion and contamination metrics of the extracted bins were estimated using CheckM [58]. The resulting bins were collectively processed to produce consolidated metagenome-assembled genomes (MAGs) using the bin_refinement module in wetaWRAP.

### Statistical analyses and OTU Classification

Statistical analyses were performed using the ‘vegan’ package [59] in R software version 4.1.1. Tests with P<0.05 were considered to be statistically significant. Shapiro-Wilk normality test was performed for each analysis. ANOVA was performed when abundance data were normally distributed. A non-parametric Kruskal-Wallis one-way analysis of variance was performed when the data were not normally distributed [60]. In parallel, to test the significance of the differences between 2 samples (i.e., between heavy and light fractions), two-tailed independent t-tests were done. For all OTU-based statistical analyses, the data set was normalized by a Hellinger transformation [61] using the decostand function. For beta-diversity, principal coordinates analysis (PCoA) ordination of Hellinger distances was carried out using the ‘pcoa’ function. Heatmaps were constructed with ‘pheatmap’ package [62] for the OTUs explaining most of the differences between samples. Principal component analysis (PCA) of the Hellinger transformed data was performed using the prcomp function. The 20 OTUs explaining most of the differences between samples were defined as the OTUs contributing the largest absolute loadings in the first and second dimensions of the PCA [60], obtained from the rotation output file. Hierarchical clustering of the distance matrix was carried out with the “ward.D2” method using ‘hclust’ function.

### Taxonomy Analysis

A representative sequence of each OTU was aligned against the SILVA 16S rRNA gene database using the naïve Bayesian classifier (bootstrap confidence threshold of 80%) by using the mothur software platform [63].

The taxonomic classification of the MAG was performed as explained previously [64]. Briefly, DNA–DNA hybridization (dDDH) was conducted using the Type Strain Genome Server (TYGS) [65]. Amino-acid comparisons between the MAG retrieved in this study and their closest relative strains were calculated based on reciprocal best hits (two-way amino acid identity AAI) using the enveomics collection [66]. Finally, a phylogenomic tree was created using the automated multi-locus species tree (autoM-LST) pipeline [67]. AutoMLST determines closely related genomes based on alignment of >90 core genes, and the closest species were determined based on percent average nucleotide identity (ANI).

### Antimicrobial resistance genes

Since only one MAG was recovered in this study, the unbinned-assembled reads (from heavy and light fractions) were also analysed. Therefore, all (MAG-1, unbinned heavy fractions and unbinned light fractions) reads were screened for antimicrobial resistance genes (ARGs) using the public database Resfinder version 4.1 [68].

## Supporting information

Hernandez-SI

## Acknowledgements

The authors thank Mike Cotton (University of Southampton) for helping with soil sampling. This study was supported by NERC pump priming grant (NE/N02026X/1). S.R. was supported by a NERC-NEOF grant (NEOF1490). M.H. gratefully acknowledges Royal Society Dorothy Hodgkin Research Fellowship (DHF\R1\211076).

## Authors’ contributions

MH, CWK and MGD planned the experiments. MH carried out experimental work. MH carried out bioinformatic analysis. MH, SR and MGD analysed the data. MH and SR wrote the manuscript. All authors read and approved the final manuscript.

## Funding

Work on this project was supported by NERC pump priming (grant number NE/N02026X/1).

## Availability of data and materials

Sequence data were deposited in the NCBI Sequence Read Archive (SRA) under accession number PRJNA428598 for 16S rRNA gene sequences, PRJNA602606 for raw metagenome data, and PRJNA778335 for MAG-1.

## Ethics approval and consent to participate

Not applicable.

## Consent for publication

Not applicable.

## Competing interests

The authors declare that they have no competing interests.

## Notes

### Competing Interest Statement

The authors have declared no competing interest.

