## Supplementary material for "Identification of diverse antibiotic resistant bacteria in agricultural soil with H_2_^18^O stable isotope probing and metagenomics": Hernandez-SI

*Running title: Multi-drug resistant bacteria in agricultural soil*

*Corresponding author:

Marc G. Dumont, University of Southampton, Life Sciences Building 85, Highfield Campus, Southampton SO17 1BJ, UK,

**Supplementary Methods**

Preliminary experiment 1: Incubation procedures

Meropenem (Sigma Aldrich, UK) at a concentration of 50 µg/ml was added to the soil (2 g) and incubated in water (10 ml) for 12 days at 180 rpm, room temperature (~22°C) and dark. Meropenem was added to the slurry incubations every three days. Samples (15 µl) were taken every day and also before and after every addition of the antibiotic. Agar diffusion test were performed by using LB Agar (tryptone 10 g/l, yeast extract 5 g/l, NaCl 5 g/l, Agar 15 g/l) media and *Escherichia coli* K12. Plates were incubated at 37°C and the presence of halo were measured after incubation.

Preliminary experiment 2: Incubation procedures for multi-drug test

An incubation with multi-drug test was performed in order to determine the suitable antibiotics to be used for further labelling experiment. Antibiotics were chosen because of their mechanism action, i.e, cell was synthesis action: cefotaxime (ctx) and meropenem (mem); protein synthesis action: gentamicin (gen) and amikacin (ami) for their 30S subunit synthesis inhibition and erythromycin (ery) for its 50S subunit synthesis inhibition; nucleid acid synthesis action: ciprofloxacin (cip) for its DNA gyrase synthesis inhibition; trimethoprim (tmp), sulfamethizole (smz) for their folate synthesis inhibition; rifampicin (rif) for its RNA polymerase synthesis inhibition. These antibiotics have been found in the genome of *Klebsiella pneumonia* [1]. Antibiotics (50 µg/ml) was added to soil (1 g) and incubated with 1 ml of water at 180 rpm, dark and room temperature for 4 days. Antibiotics were added at the beginning of the incubation and after 48 h.

Table S1. Physico-chemical parameters from Chilworth agricultural soils

| | Soil 1 | Soil 2 | Soil 3 |
|---|---|---|---|
| pH | 6.2 | 6.1 | 6.2 |
| P (mg/l) | 107 | 117 | 126 |
| K (mg/l) | 219 | 175 | 216 |
| Mg (mg/l) | 89 | 150 | 110 |
| Total N (g/kg) | 5.25 | 4.2 | 5.6 |
| dry matter % | 89.88 | 82.85 | 85.23 |
| Cu (mg/l) | 9.15 | 9.15 | 10.7 |
| Org. matter (%) | 6.5 | 9.3 | 7.4 |

Table S2: Heat map illustrating growth in all replicates (dark grey), in some of the replicates (light grey) and no growth of selected antibiotics and their combination with meropenem (mem) during a 4-day incubation of soil with water. ctx: cefotaxime, mem: meropenem; gen: gentamicin; ami: amikacin; ery: erythromycin; cip: ciprofloxacin; tmp: trimethoprim; smz: sulfamethizole; rif: rifampicin.

| Antibiotic | d-1 | d-2 | d-2* | d-3 | d-4 | Antibiotic | d-1 | d-2 | d-2* | d-3 | d-4 |
|---|---|---|---|---|---|---|---|---|---|---|---|
| ctx | | | | | | ctx+mem | | | | | |
| gen | | | | | | gen+mem | | | | | |
| ami | | | | | | ami+mem | | | | | |
| cip | | | | | | cip+mem | | | | | |
| tmp | | | | | | tmp+mem | | | | | |
| smz | | | | | | smz+mem | | | | | |
| rif | | | | | | rif+mem | | | | | |
| ery | | | | | | ery+mem | | | | | |
| mem | | | | | | all together | | | | | |

Antibiotic still present in the slurry for all 3 replicates
antibiotic still present in the slurry for at least 1 replicate
antibiotic not present in the slurry
*second addition of the antibiotic

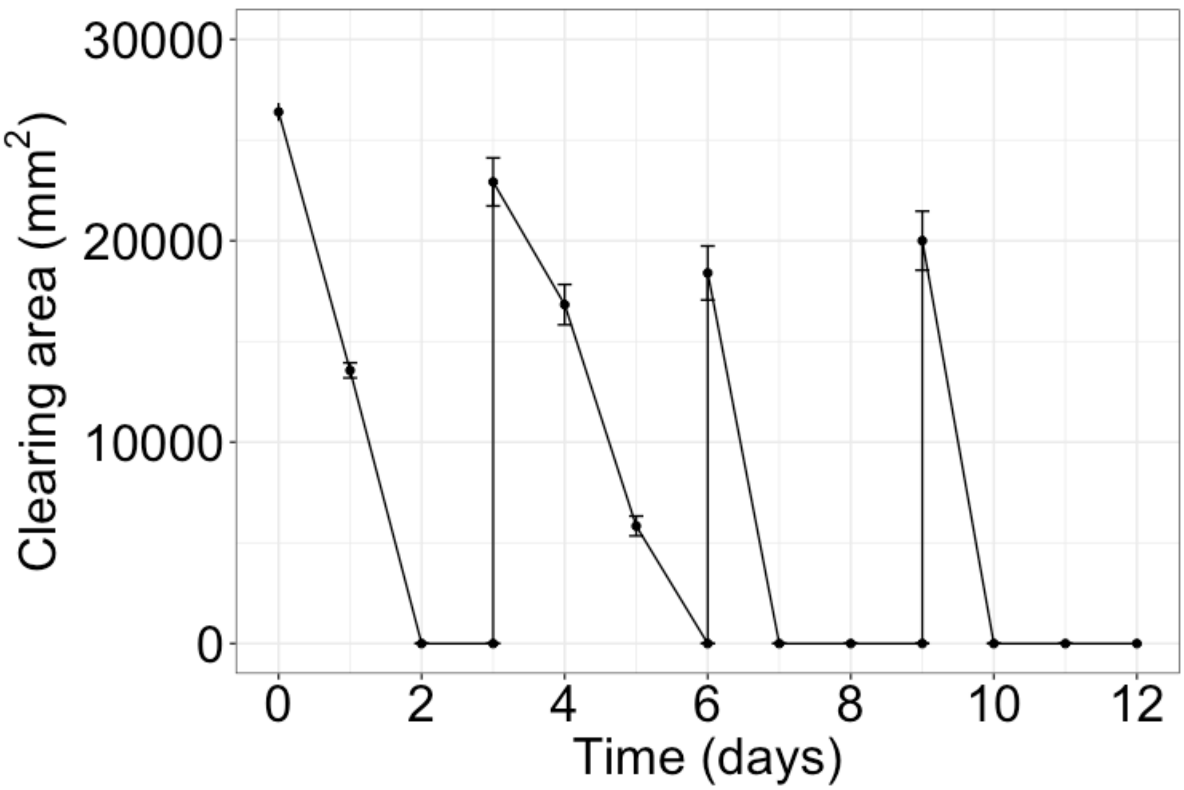


Figure S1 Degradation kinetics of meropenem during 12 days of incubation. Meropenem was added to the slurry every 2 days. Values are average and bars above the values indicate standard deviation of triplicates.

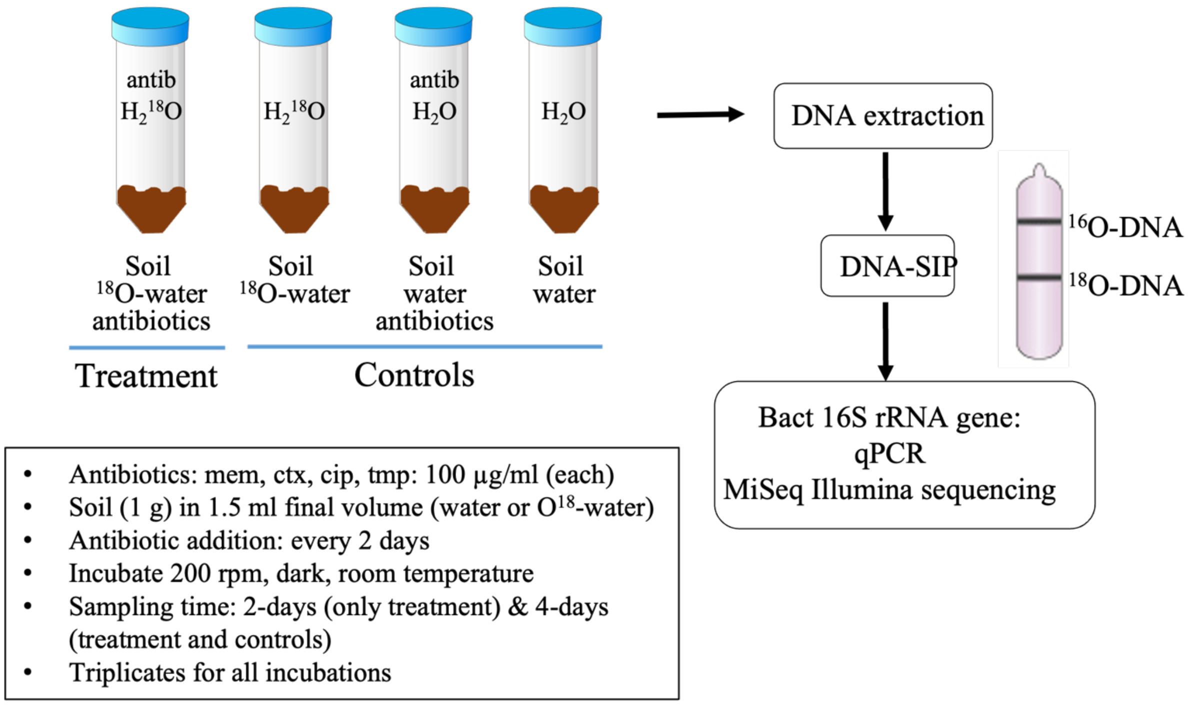


Figure S2 Scheme depicting the setting-up of the labelling incubations with antibiotics and $H_2{}^{18}O$.

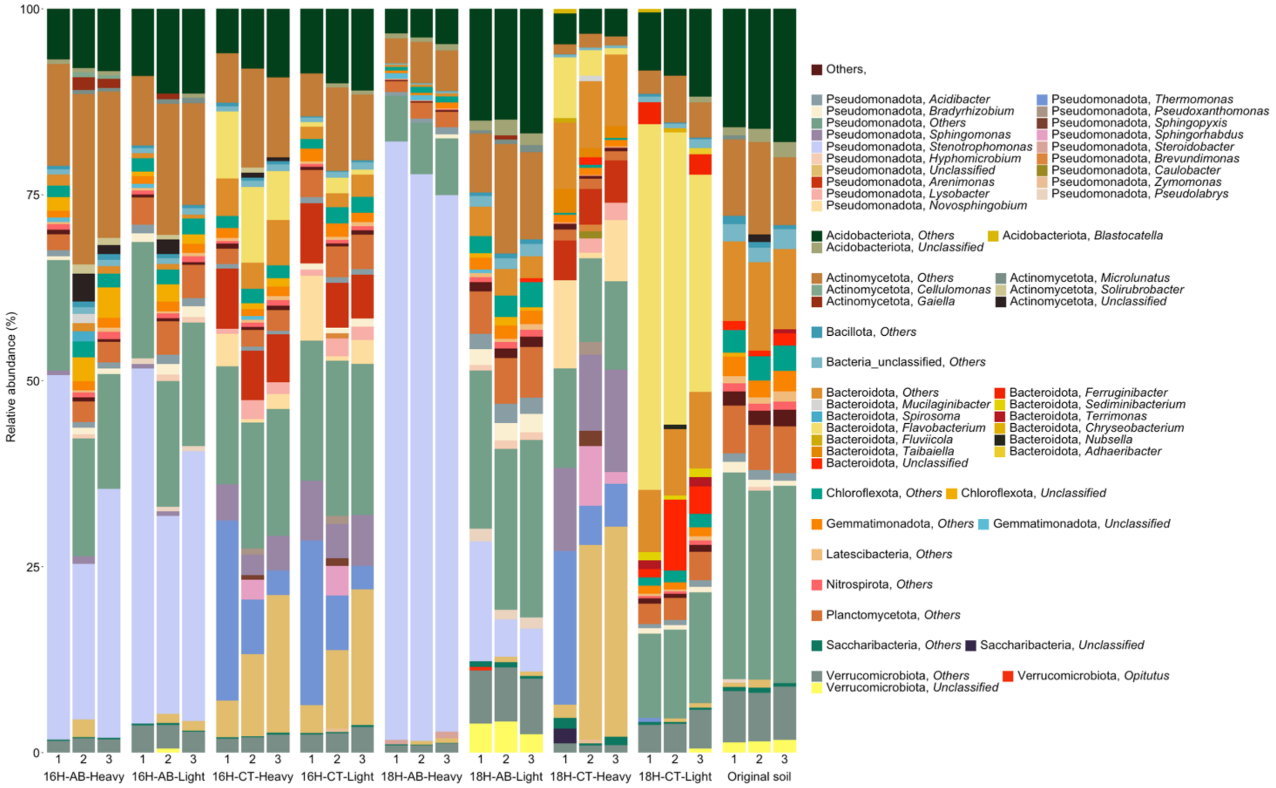


Figure S3 Relative abundance of microbial communities at the phylum level identified in the "heavy" and "light" fractions of DNA extracted from soils incubated with $H_2^{18}O$ and $H_2^{16}O$ in the presence and absence of antibiotics. 16H: incubation with $H_2^{16}O$; 18H: incubation with $H_2^{18}O$; AB: incubation with antibiotics; CT: incubation without antibiotics; Heavy: "heavy" fractions of the extracted soil DNA; Light: "light" fractions of the extracted soil DNA.

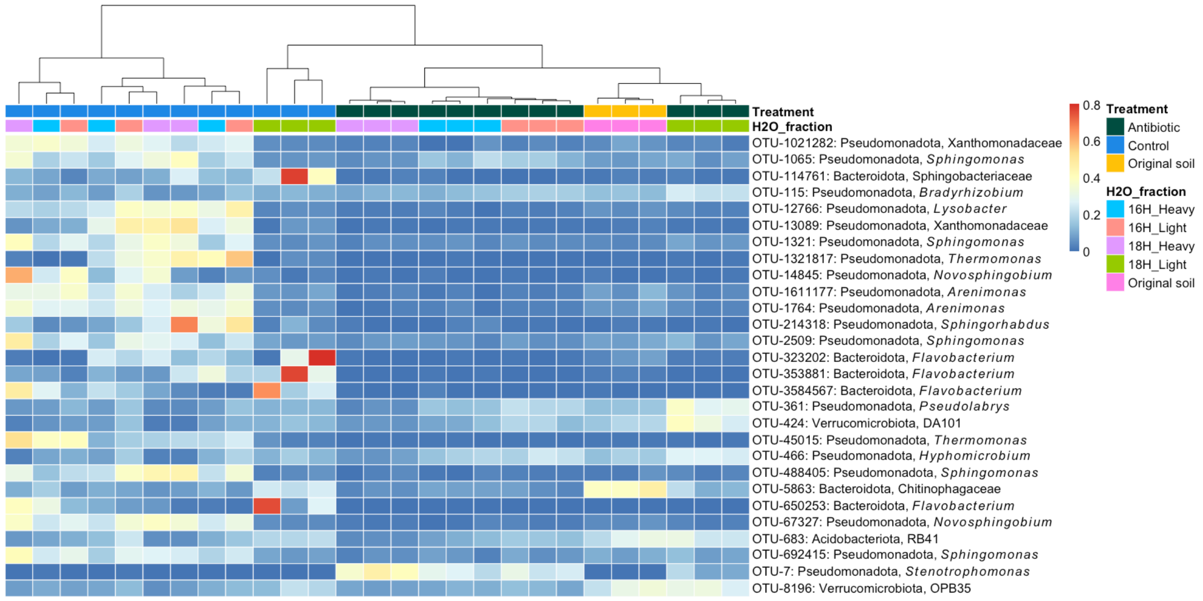


Figure S4 Heatmap of the most relevant bacterial OTUs identified in the “heavy” and “light” fractions of DNA extracted from soils incubated with $H_2{}^{18}O$ and $H_2{}^{16}O$ in the presence and absence of antibiotics. 16H: incubation with $H_2{}^{16}O$; 18H: incubation with $H_2{}^{18}O$; AB: incubation with antibiotics; CT: incubation without antibiotics; Heavy: “heavy” fractions of the extracted soil DNA; Light: “light” fractions of the extracted soil DNA.
